# Requirement of GrgA for *Chlamydia* infectious progeny production, optimal growth, and efficient plasmid maintenance

**DOI:** 10.1101/2023.08.02.551707

**Authors:** Bin Lu, Yuxuan Wang, Wurihan Wurihan, Andrew Cheng, Sydney Yeung, Joseph D. Fondell, Zhao Lai, Danny Wan, Xiang Wu, Wei Vivian Li, Huizhou Fan

## Abstract

*Chlamydia*, an obligate intracellular bacterial pathogen, has a unique developmental cycle involving the differentiation of invading elementary bodies (EBs) to noninfectious reticulate bodies (RBs), replication of RBs, and redifferentiation of RBs into progeny EBs. Progression of this cycle is regulated by three sigma factors, which direct the RNA polymerase to their respective target gene promoters. We hypothesized that the *Chlamydia-*specific transcriptional regulator GrgA, previously shown to activate σ66 and σ28, plays an essential role in chlamydial development and growth. To test this hypothesis, we applied a novel genetic tool known as dependence on plasmid-mediated expression (DOPE) to create *Chlamydia trachomatis* with conditional GrgA-deficiency. We show that GrgA-deficient *C. trachomatis* RBs have a growth rate that is approximately half of the normal rate and fail to transition into progeny EBs. In addition, GrgA-deficient *C. trachomatis* fail to maintain its virulence plasmid. Results of RNA-seq analysis indicate that GrgA promotes RB growth by optimizing tRNA synthesis and expression of nutrient-acquisition genes, while it enables RB-to-EB conversion by facilitating the expression of a histone and outer membrane proteins required for EB morphogenesis. GrgA also regulates numerous other late genes required for host cell exit and subsequent EB invasion into host cells. Importantly, GrgA stimulates the expression of σ54, the third and last sigma factor, and its activator AtoC, and thereby indirectly upregulating the expression of σ54-dependent genes. In conclusion, our work demonstrates that GrgA is a master transcriptional regulator in *Chlamydia* and plays multiple essential roles in chlamydial pathogenicity.

**IMPORTANCE:** Hallmarks of the developmental cycle of the obligate intracellular pathogenic bacterium *Chlamydia* are the primary differentiation of the infectious elementary body (EB) into the proliferative reticulate body (RB) and the secondary differentiation of RBs back into EBs. The mechanisms regulating these transitions remain unclear. In this report, we developed an effective novel strategy termed DOPE that allows for the knockdown of essential genes in *Chlamydia*. We demonstrate that GrgA, a *Chlamydia*-specific transcription factor, is essential for the secondary differentiation and optimal growth of RBs. We also show that GrgA, a chromosome-encoded regulatory protein, controls the maintenance of the chlamydial virulence plasmid. Transcriptomic analysis further indicates that GrgA functions as a critical regulator of all three sigma factors that recognize different promoter sets at developmental stages. The DOPE strategy outlined here should provide a valuable tool for future studies examining chlamydial growth, development, and pathogenicity.

## INTRODUCTION

*Chlamydia* is an obligate intracellular bacterium possessing a unique developmental cycle (1). The cycle involves two morphologically and functionally distinct cell types known as the elementary body (EB) and reticulate body (RB). The EB, approximately 0.3 µm in diameter, has DNA condensed by histones (2) and an outer membrane containing proteins crosslinked with disulfides (3). It is capable of temporary survival in extracellular environments despite having limited metabolic activities and is responsible for infecting host cells to initiate the developmental cycle. After EB is inside a host cell, its cysteine-rich outer membrane proteins undergo reduction, and DNA decondensation occurs. These processes enable EBs to differentiate into larger (up to 2.0 µm in diameter) RBs that replicate within a cytoplasmic vacuole known as an inclusion. After multiple rounds of replication, RBs asynchronously differentiate back into non-replicating EBs (1, 4). The newly formed EBs, upon release from host cells, can either infect other cells within the same host or transfer to new hosts. Unlike EBs, any released RBs are unable to initiate new developmental cycles.

The chlamydial developmental cycle is transcriptionally regulated (1, 5, 6). After EBs enter host cells, early genes are activated during the first few hours enabling primary differentiation into RBs. Starting at around 8 hours post-infection, midcycle genes, representing the vast majority of all chlamydial genes, are expressed enabling RB replication. At around 24 hours post-infection, late genes are activated to enable the secondary differentiation of RBs back into EBs.

Sigma factor is a subunit of the RNA polymerase (RNAP) holoenzyme that recognizes and binds specific DNA gene promoter elements, allowing RNAP to initiate transcription (7). *Chlamydia* encodes three different sigma factors termed σ66, σ28, and σ54 (8, 9). σ66 RNAP holoenzyme is active throughout the developmental cycle, whereas the σ28 and σ54 RNAP holoenzymes transcribe only a subset of late or mid-late genes (10–12).

GrgA is a *Chlamydia-*specific transcriptional regulator that binds both to σ66 and σ28 and activates the transcription of numerous chlamydial genes *in vitro* and *in vivo* (13–15). RNA-Seq analysis of *C. trachomatis* conditionally overexpressing GrgA, along with GrgA *in vitro* transcription assays, revealed two other transcription factor-encoding genes, *euo* and *hrcA*, as members of the GrgA regulon (13). Both *euo* and *hrcA* are transcribed during the early phase and midcycle (13). Euo acts as a repressor of chlamydial late genes (16), while HrcA regulates the expression of multiple protein chaperones (17), crucial for bacterial growth (^18^). These findings suggest GrgA plays an important regulatory role in chlamydial development.

To further determine the role of GrgA in chlamydial physiology, we attempted but failed to disrupt *grgA* through group II intron (Targetron (19)) insertional mutagenesis. Previously, the Valdivia group was also unable to generate *grgA-*null mutants using saturated chemical mutagenesis (20). Given that Targetron and chemical mutagenesis have been successfully used to disrupt numerous non-essential chlamydial genes [e.g., (21–27)], these negative results suggest that *grgA* is an essential gene for *Chlamydia* viability. In this work, we confirm that *grgA* is indeed an essential gene by using a novel genetic tool that we term DOPE (dependence on plasmid-mediated expression). Importantly, we show that GrgA is necessary for RB-to-EB differentiation and is further required for optimal RB growth and maintenance of normal copies of the virulence plasmid. These findings provide further evidence that GrgA is a major regulator of σ66, σ28, and σ54 target genes, and plays an important regulatory role in governing the chlamydial developmental cycle.

## RESULTS

### DOPE (dependence on plasmid-mediated expression) enables *grgA* disruption

Targetron is a group II intron-based insertional mutagenesis technology that has been used successfully to disrupt numerous chlamydial chromosomal genes (21–27). In an effort to knock out GrgA expression in *Chlamydia* and investigate its physiological functions, we utilized Targetron vectors containing spectinomycin-resistance gene-bearing group II introns specific for multiple *grgA* insertion sites. Whereas we did obtain spectinomycin-resistant chlamydiae, diagnostic PCR analysis indicated no insertion of the intron into *grgA* thus indicating nonspecific targeting. Taken together with the earlier unsuccessful attempts to generate *grgA*-null mutants using saturated chemical mutagenesis (20), we hypothesized that *grgA* is an essential gene and as such, not amenable to conventional mutagenesis approaches.

To circumvent this issue, we devised a strategy we term <u>d</u>ependence <u>o</u>n <u>p</u>lasmid-mediated <u>e</u>xpression (DOPE) to investigate the biological functions of essential genes in *Chlamydia*. Although disruption of essential genes in the wildtype bacterium causes lethality, transforming *Chlamydia* with a recombinant plasmid carrying the essential chromosomal gene downstream of an inducible promoter allows for the disruption of the chromosomal allele when the inducer is present in the culture medium. This method generates a strain with a disrupted essential gene, where withdrawal of the inducer will cause depletion of the gene products from the recombinant plasmid, allowing functional and mechanistic analyses of the essential gene.

We applied DOPE to study GrgA by constructing a plasmid named pGrgA-DOPE, which encodes an anhydrotetracycline (ATC)-inducible *grgA* allele (<u>p</u>lasmid-<u>e</u>ncoded inducible *<u>g</u>rgA* or peig) (sFig. 1). Compared to the native chromosomal *grgA* allele containing a Targetron-insertion site between nucleotides 67 and 68 (Fig. 1A top), the grgA allele in pGrgA-DOPE carries a His-tag sequence and four synonymous point mutations around the intron-targeting site (Fig. 1A middle), rendering peig resistant to the Targetron designed for this site. Since ATC-induced GrgA overexpression previously caused *C. trachomatis* growth inhibition (13, 28), we employed a weakened ribosomal binding site identified via a green fluorescence protein (GFP) reporter (sFig. 2A-D) in pGrgA-DOPE to drive the expression of His-tagged GrgA (His-GrgA). We also removed a region that contained potential alternative -35 and -10 promoter elements between the ATC-inducible promoter and the ribosomal binding site from pGrgA-DOPE (sFig. 2E).

**Figure 1.**
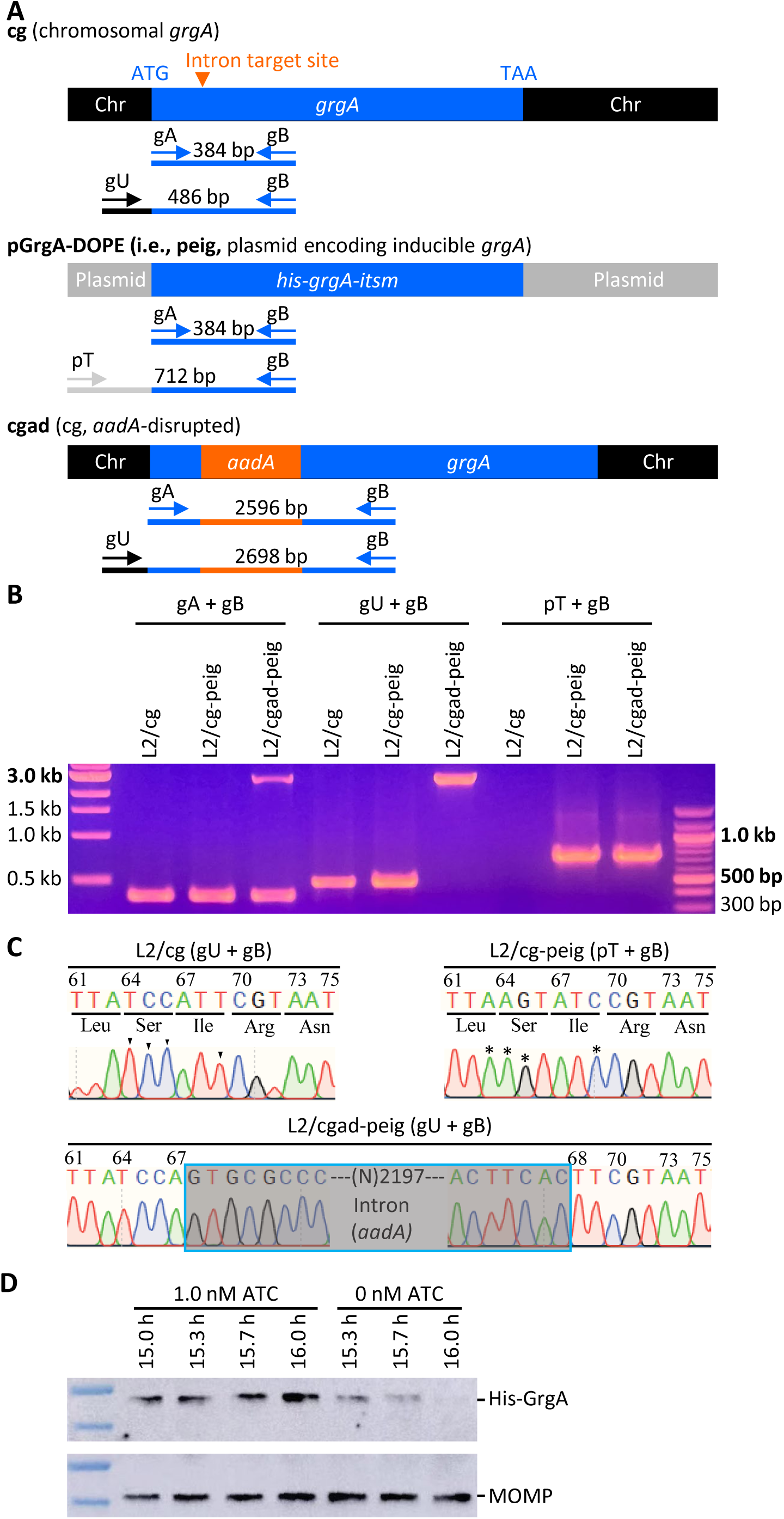
Confirmation of the disruption of the chromosome-encoded *grgA* by group II-intron and re-expression of GrgA from a transformed plasmid in the DOPE system. (A) Schematic drawings of *grgA* alleles, locations of intron-target site, diagnostic primers, and sizes of PCR products obtained with different sets of primers. Abbreviations: itsm, intron target site mutated; Chr, chromosome. (B) Gel image of PCR products amplified with DNA of wildtype *C. trachomatis* L2 with intact chromosomal *grgA* (L2/cg), L2/cg transformed with the *his-grgA-itsm* expression plasmid pGrgA-DOPE (L2/cg-peig), and L2 with *aadA-*disrupted chromosomal *grgA* complemented with pGrgA-DOPE (L2/cgad-peig) using primer sets shown in (A). (C) Sanger sequencing tracings of PCR products showing the intron-target site in L2/cg, mutations surrounding this site conferring resistance to intron targeting in peig, and *grgA-*intron joint regions in the chromosome of L2/cg-peig. Wildtye bases and corresponding mutated bases are shown with arrowheads and asterisks, respectively. (D) Western blotting showing time-dependent loss of His-GrgA in L2/cgad-peig upon ATC withdrawal. The membrane was first probed with an anti-major outer membrane protein antibody L2-5, striped, and then reprobed with an anti-GrgA antibody.

**Figure 2.**
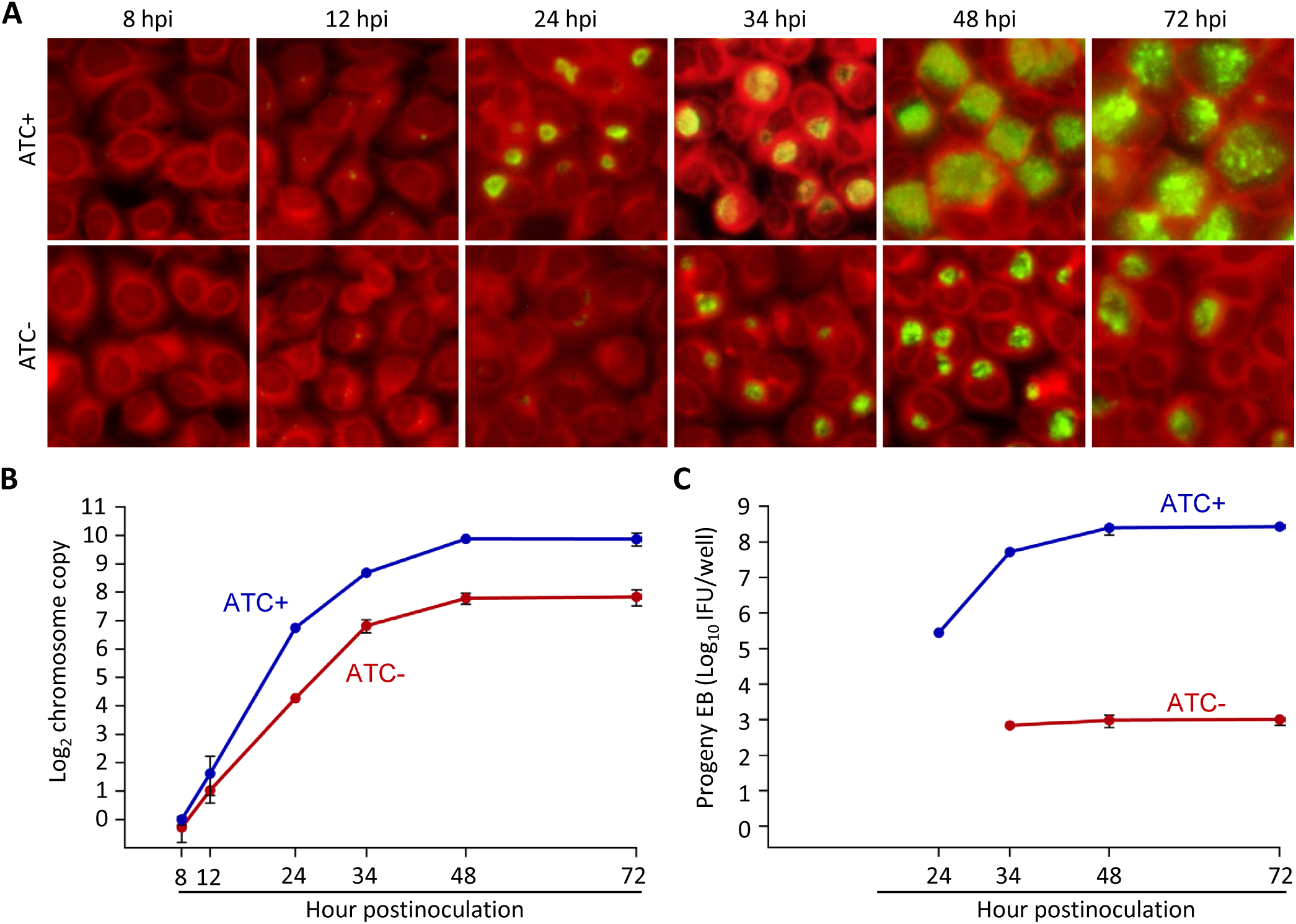
GrgA deficiency slows RB growth and disables the formation of infectious progeny. L2/cg-peig-infected HeLa cells were cultured in the presence or absence of 1 nM ATC. At indicated hpi, cultures were terminated for immunofluorescence assay (A), genome copy quantification (B), or quantification of inclusion-forming unt (C). (B, D) Data represent averages ± standard deviations of triplicate cultures.

We transformed wildtype *C. trachomatis* L2 bearing an intact <u>c</u>hromosomal *<u>g</u>rgA* (i.e., L2/cg) with pGrgA-DOPE to derive L2/cg-peig. Western blotting demonstrated comparable amounts of His-GrgA and endogenous GrgA at 12 h postinoculation (hpi) in L2/cg-peig cultures containing 0 to 5 nM ATC (sFig. 3A). We also analyzed the growth of L2/cg-peig in the presence of 0 or 1 nM ATC and found that ATC-induced GrgA overexpression did not affect the expression level of mKate2 (a red fluorescence protein encoded by pGrgA-DOPE [Fig. S1]), the inclusion size, chlamydial chromosome replication kinetics, or progeny EB production (sFig. 3B-F). These findings indicate that we can induce recombinant GrgA expression from pGrgA-DOPE at normal physiologic levels without adverse effects thereby increasing the likelihood we will be able to compensate for the loss of endogenous GrgA following the experimental disruption of the endogenous chromosomal *grgA* allele.

**Figure 3.**
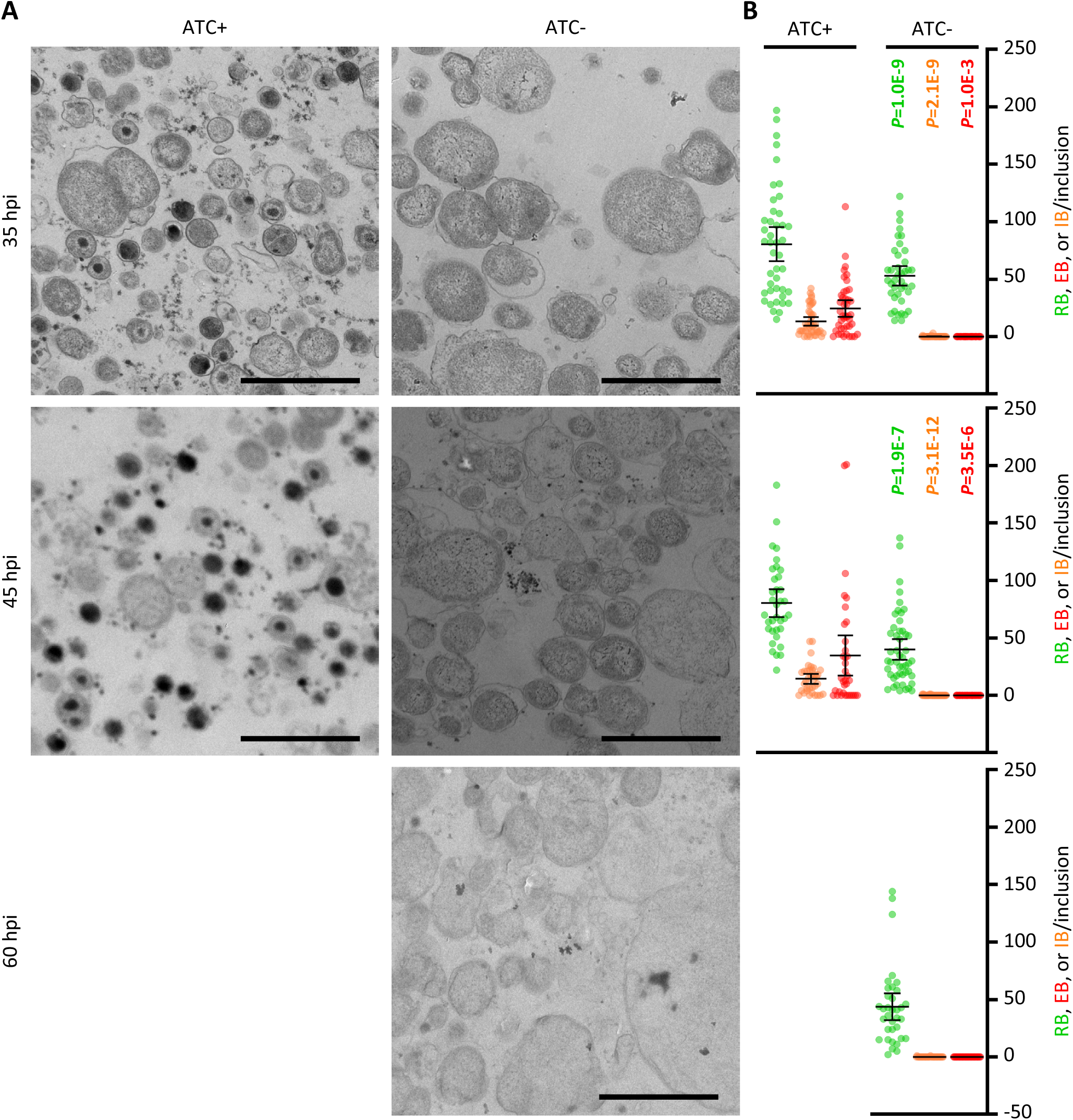
Lack of EB formation in GrgA-deficent cultures. (A) Representative electron microscopic (EM) images of L2/cgad-peig cultured in ATC-containing and ATC-free media at indicated hpi. 60 h was not processed for EM because most inclusion already burst by that point. Note that EBs are ∼400 nm diameter cellular forms with high electron density found in ATC-containing cultures. Small irregularly-shaped electron-dense particles in both ATC-containing and ATC-free cultures are glycogen particles. Cell types are color-encoded as shown in the Y-axis labels. Size bar equals 2 µm. (B) Scattergraph of RBs, EBs, and intermediate bodies (IBs) counted from multiple inclusions.

We next transformed L2/cg-peig with the aforementioned Targetron plasmid carrying an *aadA-*containing group II intron that targets the insertion site between nucleotides 67 and 68 in *grgA* (sFig. 2). ATC was employed to induce GrgA expression from pGrgA-DOPE while selecting for chromosomal mutants carrying an intron-disrupted *grgA* using spectinomycin. PCR analysis confirmed the genotypes of L2/cg, L2/cg-peig, and the plasmid-complemented, <u>c</u>hromosomal *<u>g</u>rgA-<u>a</u>adA*-<u>d</u>isrupted L2/cgad-peig (Fig. 1A, B; sTable 1). Sanger sequencing confirmed the nucleotide sequences surrounding the intron-target site in L2/cg and L2/cg-peig, and the *grgA-*intron joint regions in L2/cgad-peig (Fig. 1C). Importantly, western blotting detected a time-dependent GrgA loss in L2/cgad-peig following ATC withdrawal (Fig. 1D, sFig. 5). By 2 h post-ATC withdrawal, His-GrgA was nearly undetectable. Collectively, the data shown in Fig. 1B-D demonstrates the successful disruption of chromosomal *grgA* in L2/cgad-peig, in which expression, and thus function, of GrgA depends on ATC-induced GrgA expression from pGrgA-DOPE.

### GrgA-deficient chlamydiae display slower RB growth and fail to form progeny EBs

To examine the impact of GrgA deficiency on *C. trachomatis* growth and development, we compared the growth of L2/cgad-peig in media with and without 1 nM ATC. Interestingly, immunofluorescence imaging using a major outer membrane protein-specific antibody revealed slower growth in ATC-free cultures as indicated by significantly smaller inclusions (Fig. 2A). Quantitative PCR analysis further confirmed the slower growth in GrgA-deficient cultures as revealed by slower chromosome replication in the absence of ATC (Fig. 2B, sFig. 3). The chromosome doubling time of L2/cgad-peig in the presence of ATC was 2 hours (Fig. 2B), identical to the previously reported doubling time of wildtype L2 (13). In contrast, the doubling time of L2/cgad-peig in the absence of ATC doubled to 4 hours (Fig. 2B). Intriguingly, ATC-free cultures exhibited a drastic reduction in the production of infectious progeny EBs (Fig. 2C). On average, each ATC-containing culture produced >10^5^ inclusion-forming units (IFUs) at 24 hpi, near 10^7^ at 34 hpi, and > 10^8^ at 48 and 72 hpi (Fig. 2C). By stark contrast, ATC-free cultures produced no detectable IFUs at 24 hpi and only about 300s EBs at 34 h and thereafter (Fig. 2C). The severe decrease in EB production occurred despite chlamydial chromosome copy number in the ATC-free cultures at 34 hpi slightly surpassing that in the ATC-containing cultures at 24 hpi (Fig. 2B). Consistent with the EB quantification assays, ultra-thin section transmission electron microscopy readily detected EBs at 36 and 45 hpi in the ATC-containing cultures, whereas no EBs were detected in ATC-free cultures at 35, 45, or 60 hpi (Fig. 3A). Collectively, these results demonstrate a requirement for GrgA for optimal RB growth and the production of EBs.

### *tetR* mutations enables GrgA expression and EBs to escape in the absence of ATC

Although ATC-containing L2/cgad-peig cultures exhibited a near complete deficiency in EB formation, we were still able to detect a very low background level of EB production (Fig. 2C). We hypothesized that spontaneous mutations in the *tetR* gene and/or tetO (TetR operator) might impair the *grgA* repression in L2/cgad-peig, thereby allowing leaky GrgA expression and the production of low levels of EBs to form in the absence of ATC. To test this hypothesis, we recovered the pGrgA-DOPE plasmids from EBs formed in ATC-free cultures and expanded them in *E. coli*. Significantly, DNA sequencing revealed a single nucleotide polymorphisms (SNP) in the *tetR* gene in each of the 10 plasmids analyzed. Fig. 4 presents their locations and their effects on the 208-aa TetR protein. Two of these SNPs lead to premature termination at codons 16 or 158, while the third one induces a frameshift at codon 64. These observations indicate that lack of an authentic TetR results in the unchecked expression of wildtype GrgA in the absence of ATC. Together with findings in Figs. 3 and 4, these data further substantiate our supposition that GrgA is necessary for EB production.

**Figure 4.**
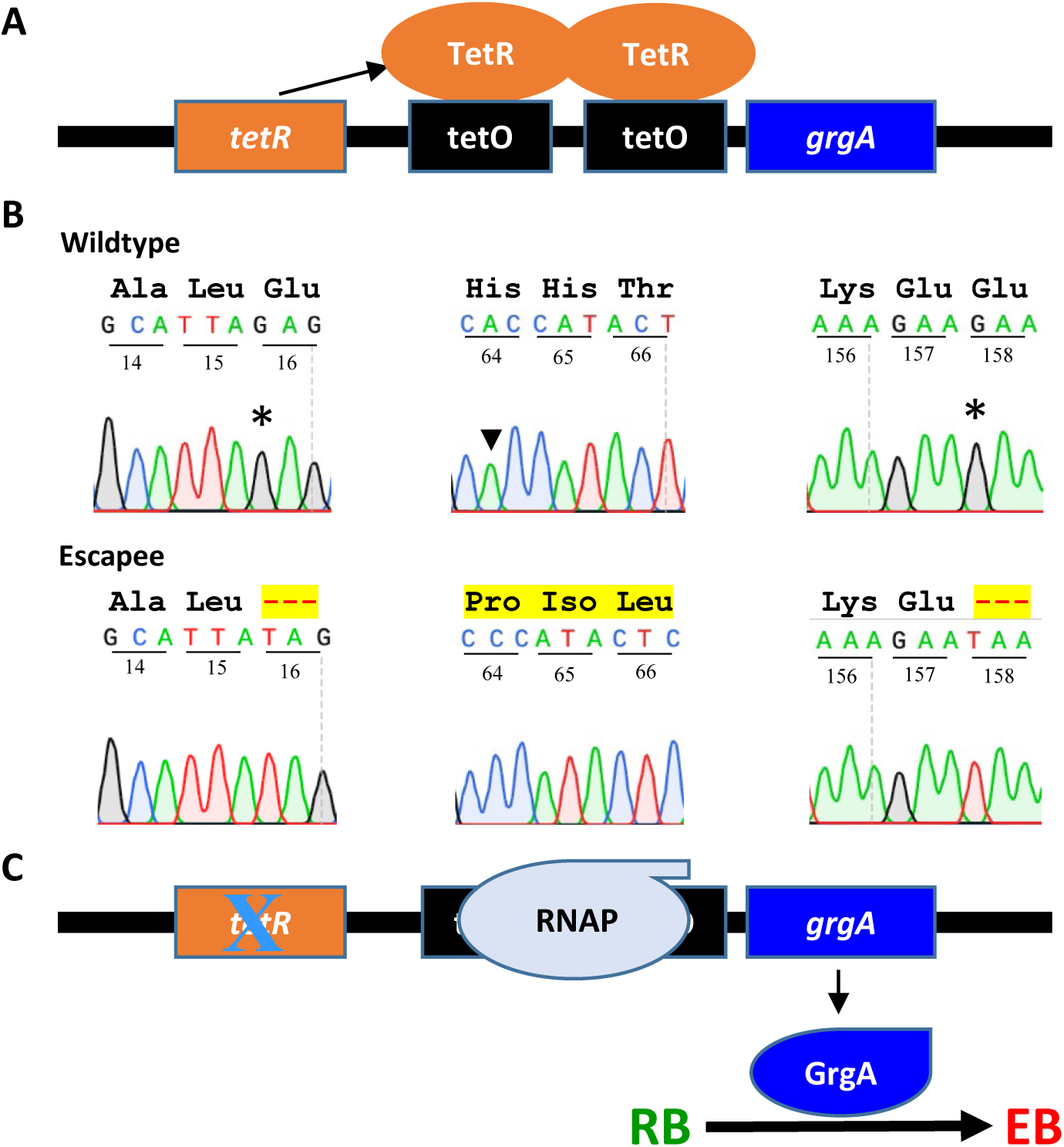
Failed *grgA* repression in pGrgA-DOPE is responsible for EB escape in ATC-free cultures of L2/cgad-peig. (A) Schematic shows mechanism for TetR-mediated *grgA* transcription repression in L2/cgad-peig cultured in the absence of ATC. (B) Mutations identified in *tetR* in pGrgA-DOPE recovered from L2/cg-peig EBs formed in the absence of ATC lead to premature translation termination or frameshift. Codon positions in *tetR* are numbered. Wild-type amino acid sequences are shown in black. Highlighted hyphens and amino acids indicate protein sequence truncation and alteration, respectively. (C) Mutations detected in (B) result in a loss in TetR-mediated *grgA* repression, leading to GrgA expression in the absence of ATC and consequent EB formation.

**Figure 5.**
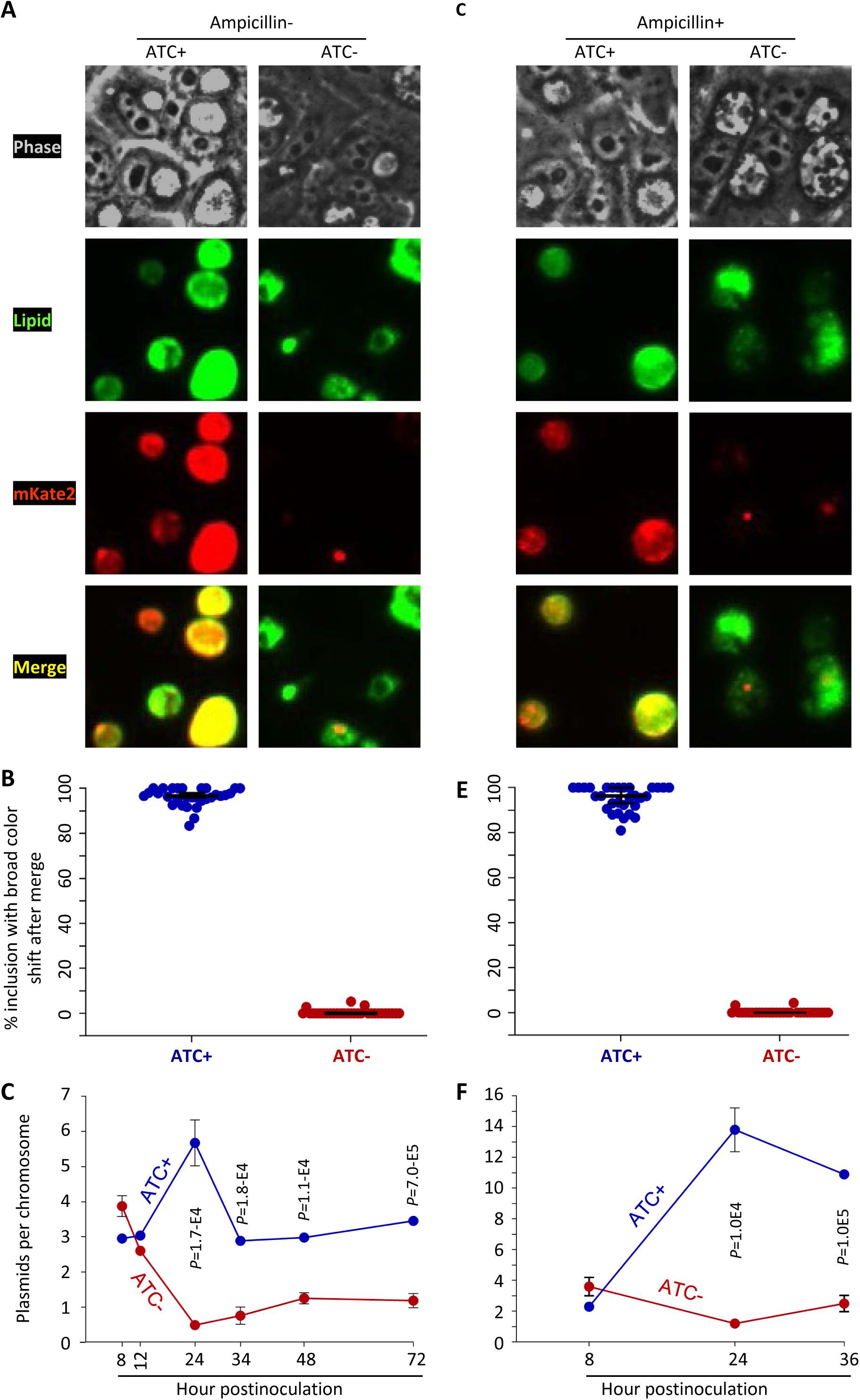
GrgA deficiency causes plasmid loss. (A) Representiative images of live cultures of L2/cgad-peig labeled with C6-NBD-ceramide (a green fluorescence lipid) in ampicillin-free media with or without 1 nM ATC. Note that the red fluorescence protein mKate2 is expressed by pGrgA-DOPE, which also encode a β-lactamase rendering ampicillin resistance. (B) Scattergraph of percentages of inclusions with broad color shift after merging green and red chanels from multiple images from cultures described in A. (C) Kinetics of plasmids per chromosome in ATC-containing and ATC-free cultures of L2/cgad-peig. Quantifiication of the pGrgA-DOPE and chromosome was carried out with qPCR as descrited in Materials and Methods. Data represent averages ± standard deviation from biological triplicates. (D) Representiative images of live cultures of L2/cgad-peig labeled with C6-NBD-ceramide in ampicillin-containing media with or without 1 nM ATC. (E) Scattergraph of percentages of inclusions with broad color shift after merging green and red chanels from multiple images from cultures described in A. (F) Kinetics of plasmids per chromosome in ATC-containing and ATC-free cultures of L2/cgad-peig. Data represent averages ± standard deviation from biological triplicates.

### GrgA-deficient chlamydiae fail to maintain the virulence plasmid

Fluorescent proteins such as mKate2 are often employed as convenient markers for tracking genetic transformants (29, 30). As depicted in sFig. 3B, L2/cg-peig produced mKate2-positive inclusions in both ATC-containing and ATC-free media. Similarly, L2/cgad-peig cultures selected in media containing ATC and spectinomycin continued to exhibit mKate2-positive inclusions. However, we were unable to identify mKate2-positive inclusions in ATC-free cultures of L2/cgad-peig, although we did notice some sporadic mKate2 signals in these cultures as detailed below.

To analyze these phenomena under higher resolution, we metabolically labeled chlamydiae with a green-fluorescing lipid (N-[7-(4-nitrobenzo-2-oxa-1,3-diazole)]) aminocaproylsphingosine (C6-NBD-ceramide) and subsequently carried out live culture fluorescence microscopy using suitable filters (31). Because L2/cgad-peig cultures exhibit slower growth in the absence of ATC, and that the number of chromosomes at 24 hpi in ATC-containing cultures is roughly equal to those at 36 hpi in ATC-free cultures (Fig. 2A, B), metabolic labeling followed by fluorescence microscopy was conducted at 24 and 34 hpi for ATC-containing and ATC-free cultures, respectively.

In ATC-containing but ampicillin-free cultures, almost 100% of inclusions exhibited strong colocalization of green fluorescence lipids and mKate2 signals, as demonstrated by significant color shifts following the merging of the lipid and mKate2 signals (Fig. 5A, B). This indicates widespread plasmid maintenance in chlamydiae. qPCR analysis detected 2.9 to 5.8 plasmids per chromosome (i.e., per cell) during the developmental cycle in ATC-enriched cultures, a count comparable to the native plasmid copy number (4.0 to 7.6 plasmids per chromosome) previously reported for wildtype *C. trachomatis* (32) (Fig. 5C). These findings suggest that L2/cgad-peig, with plasmid-expressed GrgA, is capable of maintaining its virulent plasmid even in the absence of the recombinant plasmid selection agent ampicillin (sFig. 1).

By contrast, in ATC-free cultures, no inclusions exhibiting diffused mKate2 signals of sufficient intensity to allow for significant color shifts upon merging with chlamydial lipid signals were detected (Fig. 5A and B). Indeed, the plasmid per chromosome ratio dropped to 0.5 at 24 hpi, gradually rose to 1.2 by 48 hpi, and remained at this level at 72 hpi (Fig. 5C). These observations imply that GrgA-deficient chlamydiae fail to maintain plasmid copies. Although inclusions in ATC-free cultures do not express diffused mKate2 signals, approximately 20% of the inclusions contained strong, punctate mKate2 signals, hinting at a potential plasmid segregation defect. Notably, ampicillin failed to enhance mKate2 expression (Fig. 5C, D) and only marginally increased the plasmid copy number in ATC-free cultures (Fig. 5E), suggesting that the antibiotic that typically selects for the recombinant plasmid under “normal” conditions is unable to maintain the plasmid in the absence of GrgA.

### GrgA deficiency leads to insufficient late gene activation

Given that GrgA is a transcriptional activator and that GrgA-deficient RBs fail to differentiate into EBs, we hypothesized that presumptive GrgA regulatory target genes crucial for EB formation may not be adequately activated in GrgA-deficient RBs. To identify these regulatory target genes, we performed RNA-Seq analysis on L2/cgad-peig cultured with and without ATC. We conducted RNA-Seq for ATC-containing cultures at 18 hpi and 24 hpi, corresponding to the midcycle point and early late developmental cycle point, respectively, in L2/cgad-peig (Fig. 2B, C), as well as wild type *C. trachomatis* (5, 13, 28). Considering the slower growth rate of L2/cgad-peig cultured in ATC-free medium (Figs. 3), we extended the culture time to 24 hpi and 36 hpi before extracting RNA. qPCR analysis confirmed that chromosome copy numbers in ATC-containing cultures at 18 hpi were the same as those in ATC-free cultures at 24 hpi, while chromosome copy numbers of ATC-containing cultures at 24 hpi were comparable to those of ATC-free cultures at 36 hpi (Fig. 6A).

**Figure 6.**
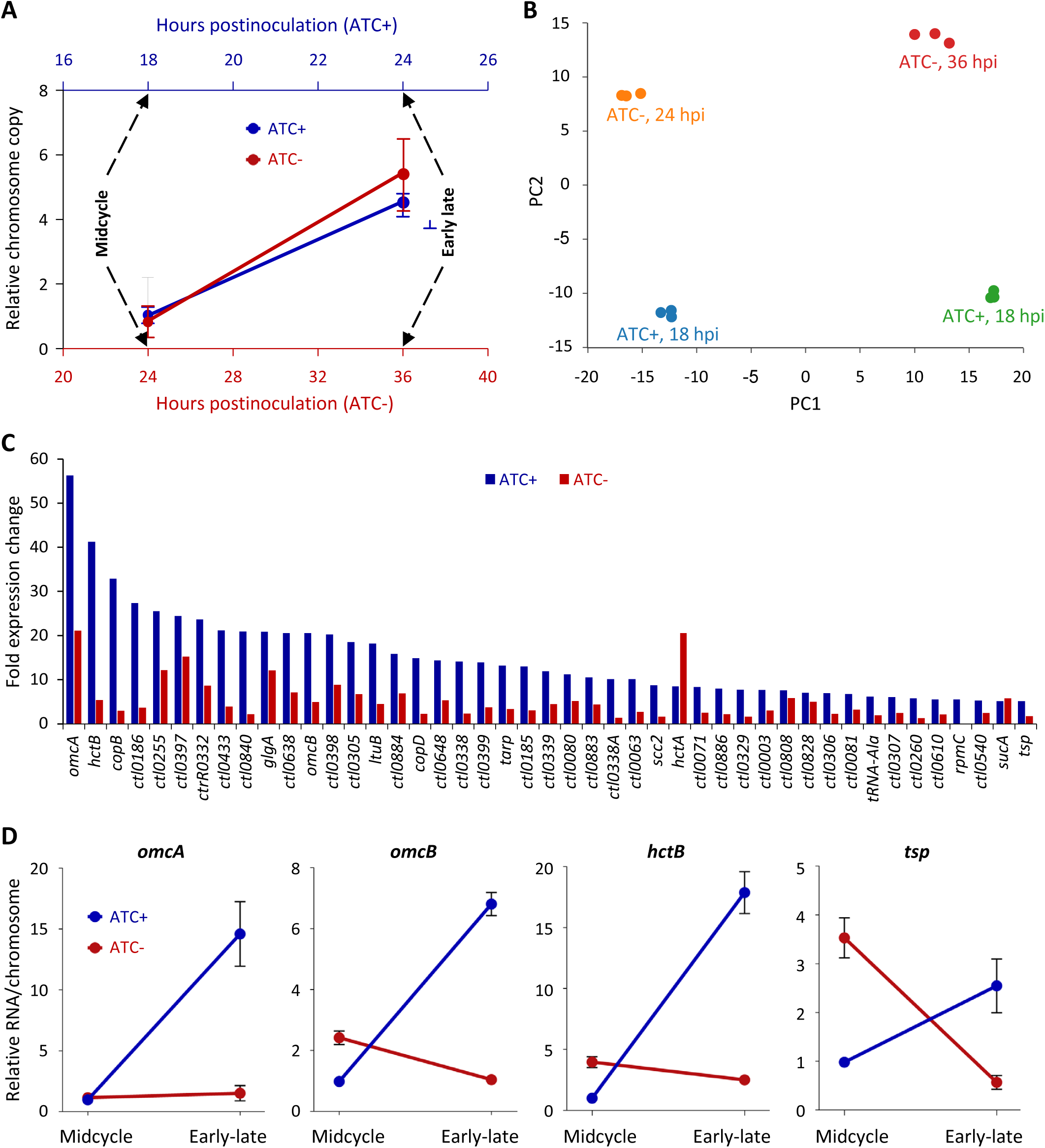
GrgA deficiency disrupts late gene activation. (A) Timing of RNA extraction for RNA-Seq analysis of ATC-containing and ATC-free cultures of L2/cgad-peig and equivalent chrosmosome copies in two types cultures at the defined midcycle points as well as the early late cycle points. (B) High intragroup consistency of RNA-Seq data revealed by pincipal component analysis. (C) Most of late genes with ≥ 5-fold increases in ATC-containing cultures had lower degree of increases in ATC-free cultures in RNA-Seq analysis. (D) Confirmation of insufficient activation of four late genes in ATC-free cultures by qRT-PCR analysis. Data were averages ± standard deviations of biological triplicates.

We obtained on average 6.3 million RNA-Seq reads mapping to the *C. trachomatis* genome per sample, representing 600 times genome coverages. Principal component analysis revealed strong consistencies within biological triplicates but notable differences among groups based on culture conditions and developmental stages (Fig. 6B). Consistent with the GrgA protein expression data in Fig. 1D and plasmid copy number data (Fig. 5), RNA-Seq detected 204- and 157-fold decreases in the transcripts of *his-grgA* encoded by pGrgA-DOPE in the ATC-free cultures of L2/cgad-peig at the corresponding midcycle and early late developmental points, respectively, compared with ATC-containing cultures (sTable 1).

We identified 155 genes activated by 2- to 56-fold (P ≤ 0.05) in ATC-containing cultures from 18 hpi (midcycle) to 24 hpi (early late stage) (sTable 2). Most likely, these late genes either drive or are consequent to the conversion of RBs into EBs. Indeed, the top two ranked genes encode an EB-enriched cysteine-rich outer membrane protein OmcA and a histone HctB, both essential for EB morphogenesis. When changes of these 155 late genes in the ATC-free cultures from 24 hpi (midcycle) to 36 hpi (early late stage) were plotted alongside the ATC-containing cultures (sTable 2), it was notable that 43 of the 45 genes with ≥ 5-fold expression increases in the ATC-containing cultures failed to increase to the same degree in the ATC-free cultures (Fig. 6C, sTable 2). For example, *hctB* increased 41.2-fold in ATC-containing cultures but only 5.3-fold in ATC-free cultures. *omcA* increased 56.2-fold in ATC-containing cultures but only 21.1-fold in ATC-free cultures (Fig. 6C, sTable 2). Similarly, *omcB* increased 20.5-fold in ATC-containing cultures but only 4.9-fold in ATC-free cultures (Fig. 6C, sTable 2). The protease-encoding gene *tsp*, thought to degrade certain RB-specific proteins towards the end of midcycle (33), exhibited a 5.5-fold increase in ATC-containing cultures but only a 1.7-fold increase in ATC-free cultures (Fig. 6C, sTable 2). Results of qRT-PCR analysis demonstrated consistent increases in *omcA, omcB, hctB,* and *tsp* in ATC-containing cultures but not ATC-free cultures from midcycle to early late cycle (Fig. 6D). These findings support the notion that GrgA is critical for adequate activation of late genes involved in the RB-to-EB conversion.

A direct comparison of RNA-Seq data from ATC-containing and ATC-free cultures of L2/cgad-peig obtained at the early late developmental point (Fig. 6A) highlighted 64 chromosomal genes that were downregulated by 2.0- to 9.1-fold in ATC-free cultures (sTable 3). As expected, the aforementioned *hctB, omcA, omcB,* and *tsp* are included in these 63 genes. Among the remaining genes, many encode T3SS structure (e.g., *copB, copD*), effector (e.g., *tarp*), or chaperone proteins (e.g., *scc2*). In late developmental stages, some T3SS effectors are secreted to enable chlamydial exit from host cells (e.g., CTL0480 and CTL0481), while other effectors are not secreted until EBs enter new host cells in the next developmental cycle (e.g., TARP) (26, 27, 34–38). These findings suggest that GrgA not only is essential for EB formation but also plays important roles in the dissemination of progeny EBs.

### GrgA deficiency downregulates expression of target genes of σ28 and σ54

Our previous *in vitro* study showed that GrgA can directly activate transcription from σ28-dependent promoters in addition to σ66-dependent promoters (14). Notably, the aforementioned *hctB* and *tsp* downregulated in ATC-free cultures of L2/cgad-peig at early late developmental points constitutes the entire σ28 regulon (39). This *in vivo* observation confirms that GrgA regulates the expression of σ28 genes in *Chlamydia*.

Two research groups have proposed partially overlapping gene sets as the σ54 regulon in *C. trachomatis* (10, 39). Soules et al. suggested 64 targets through overexpression of the σ54 activator *atoC* (*ctcC*) (10), while Hatch et al. proposed nearly 30 targets through σ54 depletion (39). Significantly, of the 64 genes exhibiting ≥ 2-fold decreases in ATC-free cultures, 42 genes, including the aforementioned *omcA, omcB, hctB*, and numerous T3SS-related genes, are σ54 targets proposed by either or both studies (Table 1). These observations suggest that GrgA also regulates the expression of σ54 genes.

**Table 1.** σ54 targets downregulated by GrgA deficiency at early late developmental points. Abbreviation, DT and IT, direct targets and indirect targets, respectively, defined by Soules et al. through RNA-Seq analysis of *atoC-*overexpressing chlamydiae, promoter sequence analysis, and transcription report assays in *E. coli* (10). T, targets defined by Hatch et al. through RNA-Seq analysis of σ54-depleted chlamydiae (39).

| Gene name | Product description | GrgA- | Soules et al. | Hatch et al. |
| --- | --- | --- | --- | --- |
| *ctl0338A* | Putative T3SS secreted protein | -9.07 | DT |  |
| *ctl0840* | Putative cytosolic protein | -8.42 | IT |  |
| *ctl0338* | Putative T3SS effector | -7.07 | DT |  |
| *copB* | T3SSs translocator CopB | -6.87 | IT | T |
| *ctl0329* | Putative outer membrane protein | -6.77 | IT |  |
| *ctl0339* | Phosphatidylcholine-hydrolyzing phospholipase D (PLD) protein | -6.57 | DT |  |
| *ctl0397* | T3SS exported protein | -6.55 | DT |  |
| *tarp* | Translocated actin-recruiting phosphoprotein | -6.30 | IT | T |
| *scc2* | T3SS chaperone | -6.29 | IT | T |
| *ctl0305* | Secreted, Pmp-like protein | -5.58 | DT |  |
| *ctl0398* | T3SS exported protein | -5.14 | DT |  |
| *copD* | T3SS translocator CopD | -5.07 | IT | T |
| *ctl0063* | T3SS effector | -5.02 | IT | T |
| *ctl0433* | Putative exported protein | -4.95 | IT |  |
| *ctl0071* | Hypothetical protein | -4.90 | DT |  |
| *glgA* | Glycogen synthase | -4.89 | IT |  |
| *ctl0307* | Putative secreted, Pmp-like protein | -4.71 | DT |  |
| *ctl0219* | Putative T3SS effector | -3.96 | DT |  |
| *ctl0220* | T3SS secreted protein | -3.61 | IT |  |
| *ctl0186* | Putative membrane protein | -3.56 | DT |  |
| *hct2* | Histone-like protein 2 | -3.36 | IT | T |
| *ctl0648* | Hypothetical protein | -3.36 | IT |  |
| *ctl0080* | Putative T3SS effector | -3.18 | DT |  |
| *ctl0306* | Secreted, Pmp-like protein | -3.10 | DT |  |
| *omcA* | Membrane protein | -3.07 | DT | T |
| *ctl0209* | Putative inclusion membrane protein | -3.02 | IT |  |
| *glgC* | Glucose-1-phosphate adenylyltransferase | -3.01 | DT | T |
| *ctl0883* | Putative T3SS effector | -2.90 | DT |  |
| *ctl0610* | Thioredoxin domain-containing protein | -2.80 | IT |  |
| *ctl0221* | T3SS secreted protein | -2.74 | IT |  |
| *ctl0185* | Putative membrane protein | -2.67 | DT |  |
| *ctl0081* | T3SS effector protein | -2.67 |  | T |
| *tsp* | Tail-specific protease | -2.66 |  | T |
| *ctl0540* | Putative inclusion membrane protein | -2.57 | IT |  |
| *ctl0003* | Putative T3SS chaperone | -2.53 | DT |  |
| *omcB* | Outer membrane protein OmcB | -2.46 | DT | T |
| *ctl0884* | Putative T3SS effector | -2.45 | DT |  |
| *ctl0255* | T3SS effector | -2.26 | DT |  |
| *ctl0260* | Inclusion membrane protein | -2.24 | DT |  |
| *ctl0021* | Hypothetical protein | -2.07 | DT |  |
| *ctl0638* | Hypothetical protein | -2.07 | IT |  |
| *ctl0466* | Inclusion membrane protein | -2.06 |  | T |

We noticed moderate (about 1.5-fold) yet significant decreases in transcripts of *rpoN* (σ54), *atoC* (*ctcC*), and *atoS* (*ctcB,* a sensor kinase gene cotranscribed with *atoC*) in ATC-free cultures of L2/cgad-peig at the early late developmental point. We performed qRT-PCR analysis and confirmed that all these three genes showed significant ≥ 2-fold decreases (Fig. 7). These observations suggest that GrgA regulates the expression of σ54 genes by controlling the expression levels of σ54 and its regulators of *atoC* (*ctcC*) and *atoS* (*ctcB*).

**Figure 7.**
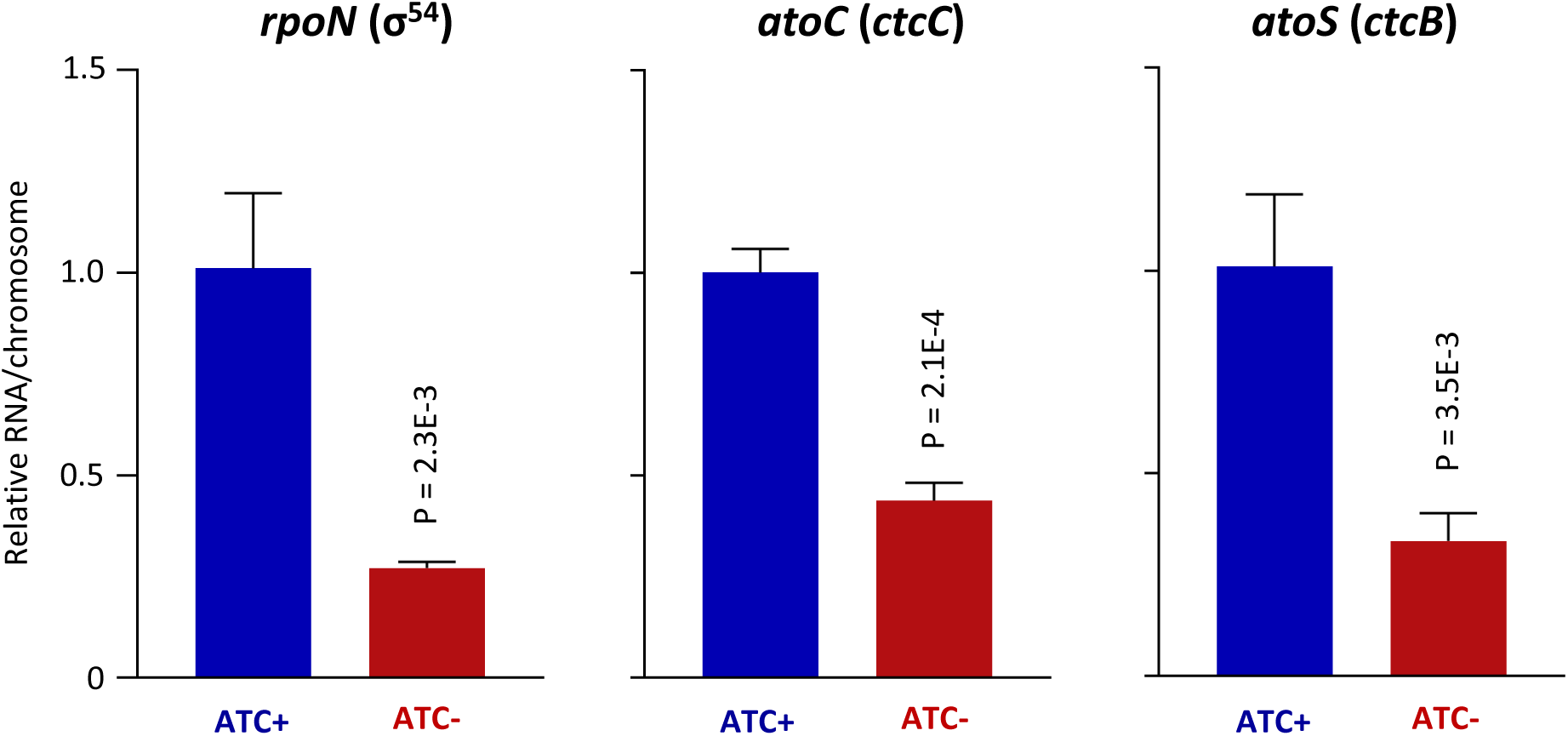
Confirmation of downregulated expression of *rpoN* and its regulators *atoC* and *ato.* Presented are qRT-PCR data (averages ± standard deviations) obtained from biological triplicates.

### GrgA deficiency disrupts the midcycle transcriptome

Our discovery of early late stage transcriptomic changes underlying EB formation deficiency prompted us to examine midcycle transcriptomic differences in L2/cgad-peig cultures with and without ATC, to further discern the mechanisms by which GrgA regulates RB growth. A total of 28 chromosomal genes exhibited a 2.0 to 4.1-fold downregulation (P < 0.05) in ATC-free L2/cgad-peig cultures as determined by RNA-Seq. Interestingly, of these 28 chromosomal genes, 7 (25%) encode tRNAs (Table 2). Of note, 10 other tRNA genes displayed 1.5- to 2.0-fold decreases (P < 0.05, sTable 1). Together, these 17 tRNAs constitute nearly half of the 37 tRNAs expressed in *C. trachomatis*, thus suggesting that GrgA supports RB growth in part, by boosting tRNA expression and protein synthesis.

**Table 2.**
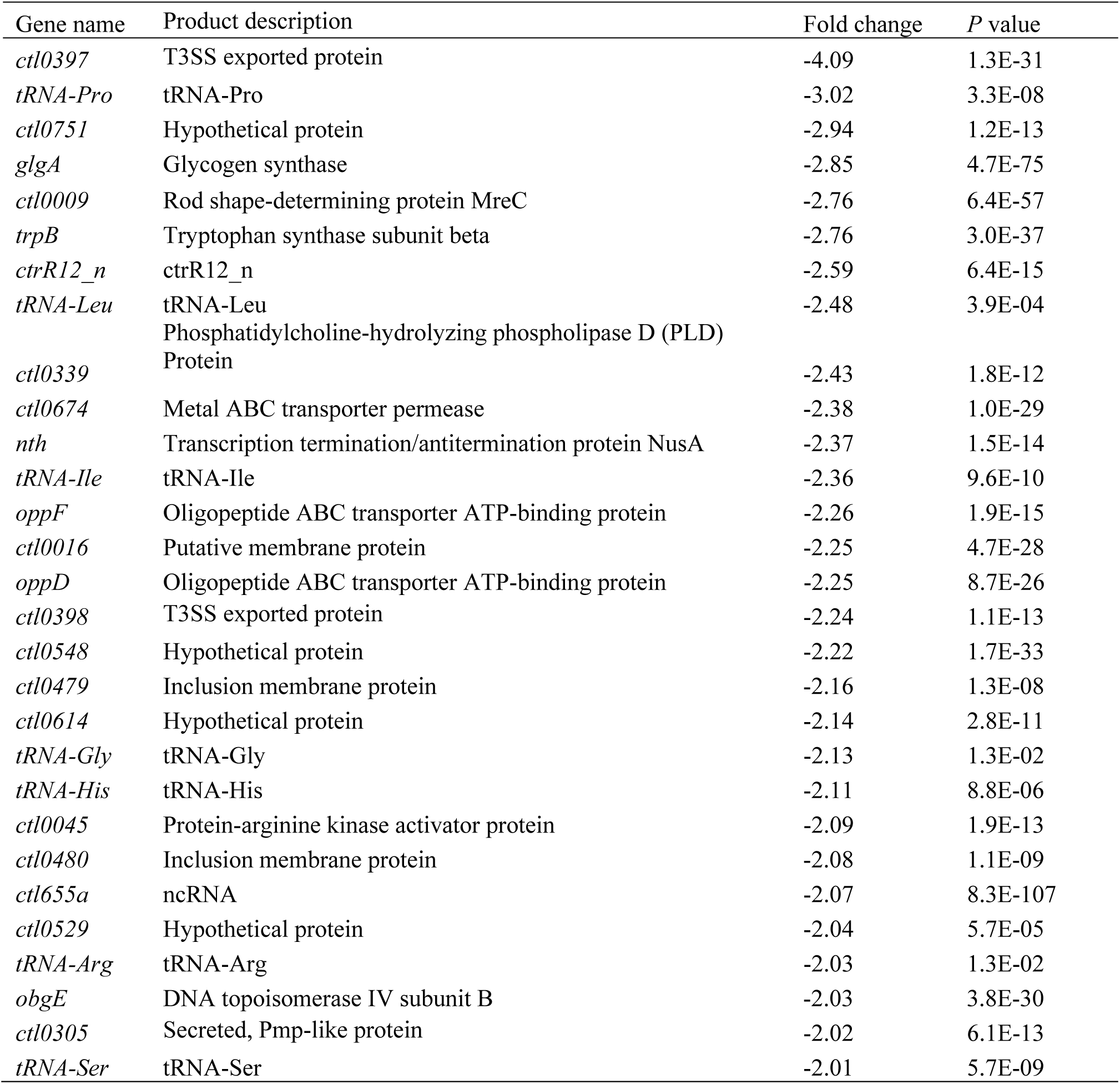
Midcycle genes downreuglated by GrgA deficiency.

Among the remaining 21 chromosomal genes with ≥ 2.0-fold expression decreases in ATC-free L2/cgad-peig cultures during the midcycle, *oppD* and *oppF* encode oligopeptide ABC transporter ATP-binding proteins, while *ctl0674* encodes a metal ABC transporter, and *trpB* encodes tryptophan synthase B. These changes suggest that GrgA promotes RB growth by upregulating both nutrient acquisition from host cells and *de novo* tryptophan biosynthesis, which both are important for chlamydial growth (40, 41).

Two additional genes of interest that are downregulated in ATC-free cultures of L2/cgad-peig during the midcycle are *obgE* and *nth*, which encode DNA topoisomerase IV subunit B and endonuclease III, and are required for DNA replication and repair, respectively. These data suggests that GrgA plays a potential role in the regulation of DNA replication and repair during RB growth.

Interestingly, 70 chromosomal genes showed 2.0- to 5.3-fold increases in ATC-free cultures at the early late stage (*P < 0.05*) compared with ATC-containing cultures (sTable 4). The number of upregulated genes is 2.5-fold higher than the number of downregulated genes discussed above (Table 1). This finding raises the possibility that GrgA also functions to regulate the silencing of large networks of developmental genes. Most notably, two well-characterized transcription factors, *euo* (late gene repressor) and *hrcA* (heat-inducible transcriptional repressor), demonstrated aberrantly increased expression in the ATC-free cultures during the midcycle (sTable 4). The increased expression of *euo* and *hrcA* was surprising because previous studies found that GrgA overexpression also increased euo and hrcA expression (13). qRT-PCR analysis confirmed that both *euo* and *hrcA* were indeed increased in ATC-cultures of L2/cgad-peig (sFig. 4). These findings strengthen the notion that GrgA serves as a master transcriptional regulator in *Chlamydia*.

Although RB replication is apparently inhibited in the absence of GrgA, we observed that four genes involved in DNA replication and repair showed increased expression in ATC-free cultures of L2/cgad-peig during the midcycle time point. These include *ruvB* (Holliday junction DNA helicase RuvB), *recA* (recombinase A), *ihfA* (DBA-binding protein Hu), and *ssb* (single-stranded DNA-binding protein). Intriguingly, all four genes are also upregulated when chlamydial growth is halted in response to heat shock (42), thus suggesting a potential regulatory role for GrgA in modulating environmental stress responses in *Chlamydia*. Similarly, four protease genes (*lon, htrA, ftsH*, and *ctl0301*) and four protein chaperone genes (*clpB, clpC, groEL*, and *groES*) also displayed increased expression in ATC-free cultures during the midcycle. In addition to the aforementioned tRNA downregulation, the upregulated protease and chaperone genes likely contribute to protein homeostatic imbalance, resulting in RB growth inhibition.

Taken together, midcycle RNA-Seq data suggest that GrgA promotes RB growth by optimizing the expression of tRNAs and certain nutrient transports. Furthermore, GrgA may function as both a transcriptional activator and a transcriptional repressor.

## DISCUSSION

### DOPE as a valuable tool for study essential gene in *Chlamydia* and other obligate intracellular organisms

Since the first demonstration of reproducible *Chlamydia* transformation using a shuttle vector 12 years ago (29), the research community has leveraged this reverse genetic tool to investigate gene function via ectopic overexpression, insertional mutagenesis, deletion, and other methods (10, 21–27, 43–47). Nonetheless, the lack of effective strategies to disrupt truly essential genes, particularly those whose overexpression is toxic, has hampered research in *Chlamydia* and other biological systems. In this study, we developed a novel, tightly regulated, inducible expression system termed DOPE, which shares similarity with a system recently reported by Cortina *et al* (46). DOPE facilitates the functional examination of an essential gene by permanently disrupting the gene in the chromosome while conditionally depleting the gene products expressed by the complementing plasmid.

The DOPE system represents a convenient and versatile tool for establishing the essentiality of a gene and at the same time, investigating the gene’s underlying functional mechanisms. An advantage of DOPE over recently developed conditional CRISPR interference systems is that gene depletion in DOPE is achieved by omitting ATC, whereas CRISPR interference relies on ATC’s presence. The adverse effects of ATC on *Chlamydia* and other bacteria have been mitigated but not entirely eradicated. In fact, it has been documented that chlamydial growth is inhibited with ATC at concentrations ≥ 20 nM (30). This makes the application of CRISPR interference to chlamydial infection in animal models challenging, as ATC concentrations in tissues and organs are not easily regulated. Furthermore, potential concerns arise about the adverse effects of ATC on the microbiota, as studies have shown that gut microbiota influences chlamydial pathogenesis (48). In addition to ATC independence, DOPE avoids off-target effects that may occur with CRISPR systems (49).

### GrgA as one of the most important regulators of chlamydial physiology

GrgA is an exclusive *Chlamydia*-specific protein with no homologs in non-chlamydial organisms, yet highly conserved among chlamydiae, including environmental chlamydiae (15). By employing the DOPE strategy, we demonstrate that GrgA plays a critical role in sustaining RB replication efficiency and is absolutely essential for RB-to-EB differentiation (Figs. 3 and 4). RNA-Seq analysis revealed numerous mechanisms through which GrgA regulates RB growth and EB formation. Perhaps chief amongst these regulatory mechanisms, during the midcycle, GrgA-deficient RBs exhibited decreased expression of numerous tRNA genes and ABC transporter genes, while displaying increased transcripts of protease genes and chaperone genes (Table 3). These findings suggest that GrgA enables optimal RB growth in part by facilitating RB protein synthesis and nutrient acquisition and by modulating post-translational protein homeostasis. Many other gene expression changes induced by GrgA deficiency (Table 1, sTable 1) likely exert additive or synergistic effects on RB growth. In addition to regulating chromosomal gene expression, our findings here show that GrgA may also regulate RB growth by enabling plasmid replication and segregation (Fig. 5). Consistent with this notion, plasmid-free *C. muridarum* has slower RB replication kinetics (50). Mechanistically, Pgp4, a plasmid-encoded proteins, has been shown to function as a transcription regulator of chromosomal genes (51, 52).

The morphological signature features of EBs are their small size and high electron density. Indeed, RB to EB conversion necessitates DNA condensation and reorganization of the chlamydial envelope. These processes involve several histones and outer membrane proteins (2, 3), respectively. Significant in this regard, our transcriptomic analysis of the early late developmental stage suggests that GrgA facilitates RB-to-EB differentiation by activating the expression of the histone gene *hctB* and late-stage outer membrane protein genes *omcA* and *omcB*. Moreover, GrgA may aid EB formation by inducing the protease gene *tsp* and modulating numerous other genes (Fig. 3C, sTables 1 and 2).

In conjunction with optimizing RB growth and enabling EB formation, our RNA-Seq analysis suggests that GrgA plays additional important roles in the chlamydial developmental cycle. For example, numerous genes encoding T3SS structure proteins, effectors, and chaperones were observed to be downregulated in GrgA deficient chlamydiae. Among the downregulated genes were the effectors CTL0480 that interacts with the host myosin phosphatase pathway and regulates chlamydial exit (27, 35), and TARP which is secreted from EBs immediately after they are taken up by host cells and interacts with host cytoskeleton protein actin to facilitate cell entry (34, 36, 37). Decreased transcripts of *ctl0480* and *tarp* in GrgA-deficient chlamydiae were observed at the early late developmental stage, thus suggesting that GrgA helps facilitate EB exit from infected cells and invasion of new host cells, both of which are required for chlamydial dissemination.

### GrgA as a crucial regulator of chlamydial plasmid maintenance

The plasmids of *C. trachomatis* and *C. muridarum* serve as important virulence factors in these two species (53–56). Our results presented in Fig. 5 indicate that GrgA plays a crucial role in *C. trachomatis* plasmid replication and/or segregation. However, the underlying mechanism is unclear. Previous studies established that *pgp1, pgp2, pgg6,* and *pgp8* are essential for the maintenance of the plasmid. Our qRT-PCR analysis and qPCR analyses demonstrated that the transcript/plasmid ratios for all *pgp* genes are actually higher in ATC-free cultures than ATC-containing cultures (data not shown). These findings suggest that plasmid loss in GrgA-deficient chlamydiae is not caused by decreased *pgp* expression. To the best of our knowledge, other than the standard DNA replication machinery components, GrgA is the first chromosome-encoded regulatory protein required for maintaining the chlamydial plasmid.

### GrgA as a master transcriptional regulator

As a component of the RNAP holoenzyme, sigma factor recognizes and binds to specific promoter sequences (7, 57, 58). Based on the temporal expression profiles of chlamydial sigma factors and recent studies, it is generally recognized that σ66 is the principal housekeeping sigma factor, whereas σ28 and σ54 are required for the expression of certain late genes regulating the differentiation of RBs into EBs (39, 59). While controversies exist regarding the exact composition of the σ28 and σ54 regulons in *C. trachomatis* (10, 11, 39, 60), numerous previously reported σ28 and σ54 target genes were found to be downregulated by GrgA deficiency (Table 1). These findings indicate that GrgA not only regulates σ66 genes, but also σ28 and σ54 genes.

Previous studies suggest that distinct GrgA domains directly bind to σ66 and σ28 to regulate the expression of their respective target genes (14, 15) (Fig. 8). Among the numerous σ66 target genes are *rpoN* (σ54), *atoC* (activator of σ54), and *atoS* (presumed positive regulator of AtoC). Findings from this study (Fig. 7) indicate that GrgA stimulates the expression of all these three genes to upregulate σ54 target genes (Fig. 8).

**Figure 8.**
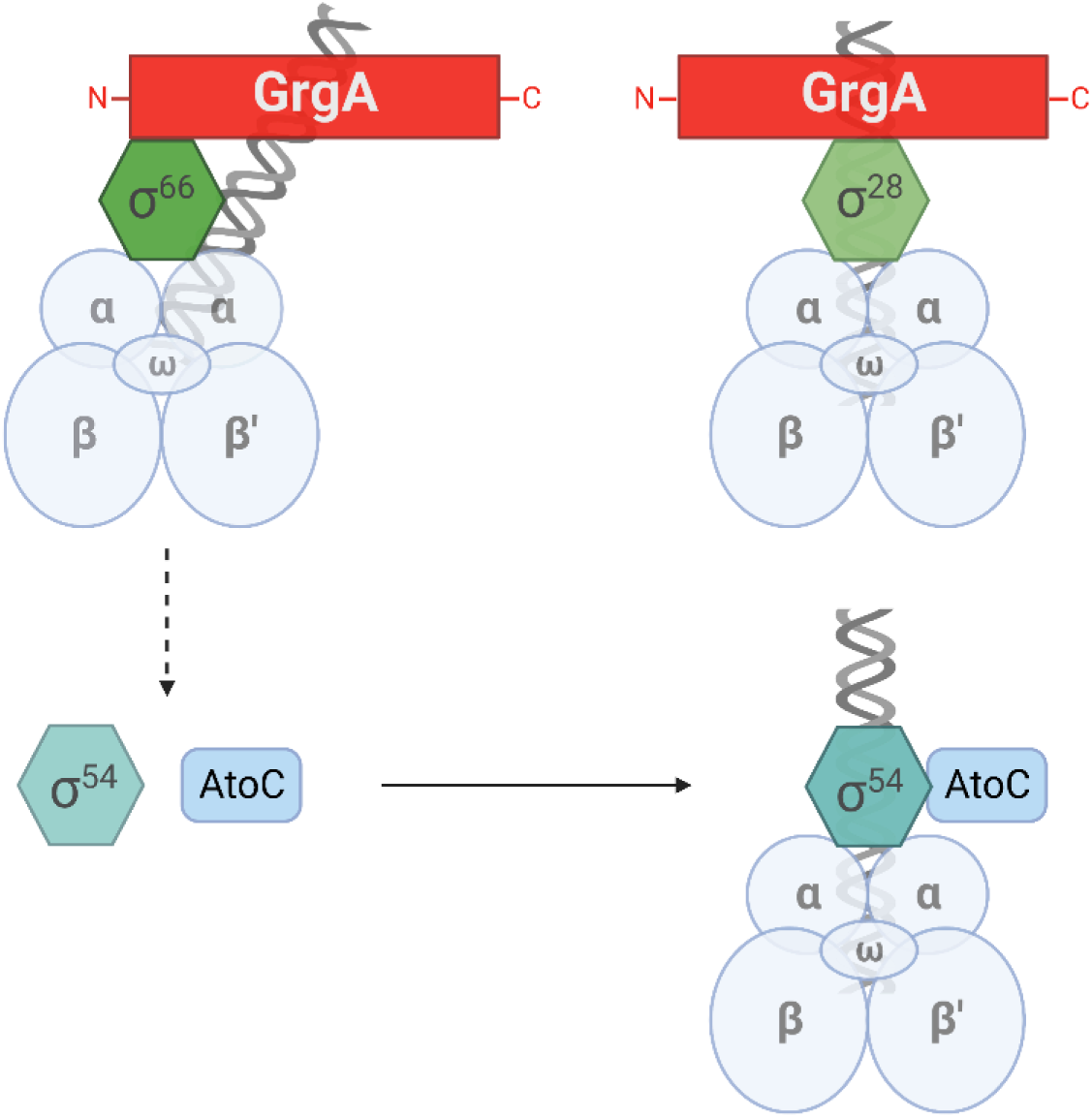
Proposed mechanisms for regulation of σ66, σ28, and σ54 target genes by GrgA. Distinct regions of GrgA interact with σ66 and σ28 to directly regulate the transcription from their target gene promoters by the RNA polymerase core enzyme (comprised of α, β, β’, and ω subunits). Among σ66-dependent genes are *rpoN* (σ54) and *atoC,* which are upregulated by GrgA. Accordingly, σ54 target genes are indirectly upregulated by GrgA. Figure was generated using paid subscription to Biorender.

*In vitro* transcription analysis established that GrgA functions as a transcriptional activator (13–15). Surprisingly, *euo* and *hrcA,* whose promoter activities were stimulated by GrgA *in vitro* and whose transcripts were increased by GrgA overexpression, exhibited increased transcripts in GrgA-deficient chlamydiae (sTables 1, 4; sFig. 4). This seemingly inconsistency raises the possibility that increased *euo* and *hrcA* expression in GrgA-deficient chlamydiae could be the result of indirect regulation. However, the fact that GrgA-deficiency resulted in 2.5 times more increased genes (sTable 4) than decreased genes (sTable 3) during the midcycle raises an alternative possibility that GrgA could function as both an activator and a repressor. It is worth noting that other bacterial transcription factors can both activate and repress genes (61–63). For example, the cyclic AMP receptor protein (CRP) binds different region of an outer membrane protein promoter and activates transcription by directly interacting with RNAP but can also repress the transcription after recruiting a transcriptional corepressor (61). We speculate that GrgA could regulate its target genes in an analogous manner. Taken together, GrgA regulates the expression of target genes of all three sigma factors, albeit through distinct mechanisms; GrgA also regulates the expression of other transcription factors. These functions validate GrgA’s role as a master transcriptional regulator in *Chlamydia*.

In summary, DOPE has enabled us to disrupt an essential chromosomal gene *grgA*. We show that GrgA serves as a checkpoint for chlamydial secondary differentiation and a crucial regulator of RB growth and plasmid maintenance. The Formation of EBs is absolutely required for dissemination of chlamydial infection within the infected host and transmission to new hosts. Because RBs and EBs share most of the immunodominant antigens (e.g., major outer membrane protein), conditional GrgA-deficient, “maturation”-defective chlamydiae are potential candidates for life attenuated *Chlamydia* vaccines, provided that strategies are in place to fully prevent EBs from escaping the gene expression regulatory system in DOPE plasmid. To the minimum, the maturation-defective chlamydiae will serve as a useful system for studying the roles of RBs in antichlamydial immunity.

## MATERIALS AND METHODS

### Vectors

pTRL2-grgA-67m (sFig. 1), which carried a *grgA* allele with resistance to intron insertion between nucleotides 67 and 68, was constructed by assembling 3 DNA fragments using the NEBuilder HiFi DNA assembly kit (New England Biolabs). All 3 fragments were amplified from pTRL2-His-GrgA (28) using Q5 DNA polymerase (New England Biolabs). Fragment 1 was generated using primers pgp3-pgp4-F and His-RBS-R (sTable 5). Fragment 2 was generated using primers RBS-His-F and GrgA-67-R (sTable 5). Fragment 3 was generated using primers GrgA-67-F and pgp4-pgp3-R (sTable 5).

pDFTT3(aadA), a Targetron vector for disrupting chlamydial genes through group II intron insertional mutagenesis (64), was a generous gift from Dr. Derek Fisher (Southern Illinois University, IL). To construct pDFTT3(aadA)-GrgA-67 (sFig. 2), which was designed for disrupting the open reading frame of *grgA*, two PCR fragments were first generated using pDFTT3(aadA) as the template. Fragment 1 was obtained using primers GrgA67_IBS1/2 and the Universal primer (sTable 5), while fragment 2 was obtained using primers GrgA67_EBS2 and GrgA67_EBS1/delta (sTable 5). The two fragments were combined and subject to PCR extension. The resulting full-length intron-targeting fragment was digested with *Hind*III and *BsrG*I and subject to ligation with *Hind*III- and *BsrG*I-digested pDFTT3(aadA). The ligation product was transformed into *E. coli* DH5α, which was plated onto LB agar plates containing 500 µg/ml spectinomycin and 25 µg chloramphenicol. The authenticity of the insert in pDFTT3(aadA)-grgA-67m was confirmed using Sanger sequencing, a service provided by Quintara Biosciences.

### Host cells and culture conditions

Mouse fibroblast L929 cells were used as the host cells for *C. trachomatis* transformation and preparation of highly purified EBs. Human vaginal carcinoma HeLa cells were used for experiments determining the effects of GrgA depletion on chlamydial growth and development. Both L929 and HeLa cell lines were maintained as monolayer cultures using Dulbecco’s modified Eagle’s medium (DMEM) (Sigma Millipore) containing 5% and 10% fetal bovine serum (vol/vol), respectively. Gentamicin (final concentration: 20 µg/mL) was used for maintenance of uninfected cells and was replaced with penicillin (10 units/mL) and/or spectinomycin (500 µg/mL) as detailed below. 37 ℃, 5% CO_2_ incubators were used for culture uninfected and infected cells.

### Chlamydiae

Wildtype *C. trachomatis* L2 434/BU (L2) was purchased from ATCC. L2/cg-peig was derived by transforming L2 EBs with pTRL2-grgA-67m using calcium phosphate as previously described(28). The transformation was inoculated onto L929 monolayer cells and selected with penicillin as previously described (28). L2/cgad-peig was derived by transforming L2/cg-peig with pDFTT3(aadA)-grgA-67m in the same manner. ATC was added to the cultures immediately after transformation to induce the expression of GrgA from pDFTT3(aadA)-grgA-67m. 12 hours later, spectinomycin D (final concentration: 500 µg/ml) was added to the culture medium to initiate selection (28). L2/cgad-peig EBs were amplified using L929 cells and purified with ultracentrifugation through MD76 density gradients (65). Purified EBs were resuspended in sucrose-phosphate-glutamate (SPG) buffer; small aliquots were made and stored at -80 ℃. We added cycloheximide to all chlamydial cultures (final cycloheximide concentration in media: 1 µg/mL) to optimize chlamydia growth. Triplicate cultures on 12-well plates were used for experiments analyzing chlamydial chromosomal and plasmid replication and EB formation.

### Immunofluorescence staining

Near-confluent HeLa monolayers grown on 6-well plates were inoculated with L2/cgad-peig at MOI of 0.3 inclusion-forming units. Following 20 min centrifugation at 900 g, cells were cultured at 37 ℃ in media containing either 0 or 1 nM ATC for 30 h. The infected cells were then fixed with cold methanol, blocked with 10% fetal bovine serum prepared in phosphate-buffered saline (PBS), and stained successively with the monoclonal L21-5 anti-major outer membrane protein antibody (66) and an FITC-conjugated rabbit anti-mouse antibody cells. Immunostained cells were finally counter-stained with 0.01% Evans blue (in PBS). Red (Evan blue) and green (MOMP) fluorescence images were acquired on an Olympus IX51 fluorescence microscope using a constant exposure time for each channel. Image overlay was performed using the PictureFrame software. The Java-based ImageJ software was then used to process the image (28).

### IFU assays

Frozen purified L2/cgad-peig EB stock or crude harvests of L2/cgad-peig cultured with or without ATC were thawed, 1-to-10 serially diluted, and inoculated onto L929 monolayers grown on 96-well plates using medium containing 1 nM ATC and 1 µg/mL cycloheximide. Following 20 min centrifugation at 900 g, cells were cultured at 37 ℃ for 30 h. Cell fixation and antibody reactions were performed as described above. Immunostained inclusions were counted under the fluorescence microscope without Evan blue counter staining.

### Diagnostic PCR and DNA sequencing

For confirming and sequencing *grgA* alleles in the chromosome and plasmid, total DNA was extracted from ∼ 1000 infected cells using the Quick-gDNA MiniPrep kit (Sigma Millipore) following manufacturer’s instructions. The resulting DNA was used as template for PCR amplification using Taq DNA polymerase (Genscript). DNA fragments resolved with electrophoresis of 1.2% Agarose gel purified using the Gel Extraction Kit (Qiagen) and subject to Sanger sequencing at Quintara Biosciences.

### Quantification of chromosome and plasmid copy numbers

To quantify chromosome copy numbers in cultures, infected cells were detached from 12-well plates using Cell Lifters (Corning). Cells and media were collected into Eppendorf tubes, centrifuged at 20,000 g at 4 ℃. The supernatant was carefully aspirated. 100 µL alkaline lysis buffer (100 mM NaOH and 0.2 mM EDTA) was added into each tube to dissolve the cell pellets. Tubes were heated at 95 ℃ for 15 min and then placed on ice. 350 µL of H_2_O and 50 µl of 200 mM Tri-HCl (pH 7.2) were added into each tube and mixed. The neutralized extracts were used for qPCR analysis (1 µL/reaction) directly for samples collected up to 24 hpi or after a 100-fold dilution for samples collected thereafter. A pair of *ctl0631* primers (sTable 5) were used for qPCR analysis to quantify chromosome copy numbers, while a pair of *pgp1* primers (sTable 5) were used to quantify plasmid copy numbers. qPCR analysis was performed with biological triplicates and technical duplicates using QuantStudio 5 real-time PCR System and Power SYBR Green PCR Master Mix (Thermo Fisher Bioscientific) (13, 28).

We took advantage of the pGrgA-DOPE plasmid to quantify the plasmid-to-chromosome ratio without preparing chlamydial chromosomal DNA free of host DNA contamination. Briefly, a pair of *grg*A primers and the aforementioned *pgp1* primers were simultaneously used to quantify pGrgA-DOPE prepared from *E. coli*, while the *grgA* primers (sTable 5) and the aforementioned *ctl0631* primers were simultaneously used to quantify chromosome of wildtype *C. trachomatis.* These analyses showed that the *grgA* primers and *pgp1* primers have the same amplification efficiencies for quantifying pGrgA-DOPE, while the *grgA* primers is 30% more efficient than the *ctl0631* primers in amplifying the chromosome. To determine plasmid-to-chromosome ratio in L2/cgad-peig, we ran qPCR analysis for *pgp1* and *ctl0631* simultaneously. We calculated plasmid-to-chromosome ratios by correcting the *ctl0631* amplification efficiency with *grgA* amplification efficiency.

### Western blotting

Detection of MOMP and GrgA was performed as previously described(28). L929 cells grown on 6-well plates were infected with L2/cgad-peig and cultured with medium containing 1 nM ATC. At 15 h, 15 h 20 min, 15 h 40 min postinoculation, cells in selected wells were switched to ATC-free medium after 3 washes. At 16 hpi, cells in each well were harvested in 200 μL of 1X SDS-PAGE sample buffer, heated at 95 ℃ for 5 min, and sonicated for 1 min (5-s on/5-s off) at 35% amplitude. Proteins were resolved in 10% SDS-PAGE gels and thereafter transferred onto PVDF membranes. The membrane was propped with the monoclonal mouse anti-MOMP MC22 antibody(67), stripped and reprobed with a polyclonal mouse anti-GrgA antibody(28).

### Transmission electron microscopy

To visualize intracellular chlamydiae up to 36 hpi, L929 cell monolayers grown on 6-well plates were infected as described above and cultured with medium supplemented with or without 1 nM ATC. For cultures up to 36 h, cells were removed from the plastic surface using trypsin, collected in PBS containing 10% fetal bovine serum, and centrifuged for 10 min at 500 g. Pelleted cells were resuspended in EM fixation buffer (2.5% glutaraldehyde, 4% paraformaldehyde, 0.1 M cacodylate buffer) at RT, allowed to incubate for 2 h, and stored at 4 ℃ overnight. To visualize intracellular chlamydiae at 45 and 60 hpi, the above procedures resulted in lysis of infected cells and inclusions. To overcome this problem, cells grown on glass coverslips were infected with and fixed without trypsinization. To prepare samples for imaging, cells were first rinsed in 0.1 M cacodylate buffer, dehydrated in a graded series of ethanol, and then embedded in Eponate 812 resin at 68 ℃ overnight. 90 nm thin sections were cut on a Leica UC6 microtome and picked up on a copper grid. Grids were stained with Uranyl acetate followed by Lead Citrate. TIFF images were acquired on a Philips CM12 electron microscope at 80 kV using an AMT XR111 digital camera. EBs, RBs, and IBs were enumerated.

### RNA isolation

Total host and chlamydial RNA were isolated from L2/cgad-peig-infected HeLa cells using TRI reagent (Millipore Sigma). DNA decontamination was achieved by two cycles of DNase I-XT (New England Biolabs) digestion. Complete removal of genomic DNA was confirmed by PCR analysis. RNA concentration was determined using Qubit RNA assay kits (ThermoFisher). Aliquots of the DNA-free RNA samples were stored at -80 °C.

### RNA sequencing and analyses

RNA-seq was performed as described with minor modifications (13, 42). Briefly, total RNA integrity was determined using Fragment Analyzer (Agilent) prior to RNA-seq library preparation. Illumina MRZE706 Ribo-Zero Gold Epidemiology rRNA Removal kit was used to remove mouse and chlamydial rRNAs. Oligo(dT) beads were used to remove mouse mRNA. RNA-seq libraries were prepared using Illumina TruSeq stranded mRNA-seq sample preparation protocol, subjected to quantification process, pooled for cBot amplification, and sequenced with Illumina Novoseq platform with 100 bp pair-end sequencing module. Short read sequences were first aligned to the CtL2 434/Bu genome including the chromosome (GCF_000068585.1_ASM6858v1), the pL2 plasmid (AM886278), and four genes cloned into the plasmid (*ha-grgA*, *bla*, *tetR* and the mKate gene) using STAR version 2.7.5a and then quantified for gene expression by HTSeq to obtain raw read counts per gene, and then converted to FPKM (Fragment Per Kilobase of gene length per Million reads of the library) (68–70). DESeq2, an R package commonly used for analysis of data from RNA-Seq studies and test for differential expression (71), was used to normalize data and find group-pairwise differential gene expression based on three criteria: P < 0.05, average FPKM > 1, and fold change ≥ 1.

### Quantitative reverse transcription-PCR (qRT-PCR) analysis

qRT-PCR was performed using QuantStudio 5 real-time PCR System (Thermo Fisher Bioscientific) and Luna Universal one-step qRT-PCR kit (New England BioLabs) as previously described (13, 28). Refer to sTable 5 for information for qRT-PCR primers.

### Data availability

RNA-seq data have been deposited into the NCBI Gene Expression Omnibus under accession number GSE234589 (reviewers’ token: qhkvogqgnpwrlsn).

## ACKNOWLEDGEMENTS

We thank Rajesh Patel (Rutgers – RWJMS EM Core Facility) for processing electron microscopy samples, Dr. Derek Fisher (Southern Illinois University) for pDFTT3(aadA), Dr. Guangming Zhong (University of Texas Health Sciences Center San Antonio for the polyclonal anti-GrgA antibody, Drs. Harlan D. Caldwell (National Institute of Allergy and Infectious Diseases) for the monoclonal anti-MOMP antibody. We also acknowledge Tara Kehair’s participation in the analysis of electron microscopic images. This work was supported by grants from the National Institutes of Health (Grant # AI140167 and AI154305 to HF and GM142702 to VWL). RNA-Seq data were generated in the Genome Sequencing Facility, which is supported by UT Health San Antonio, NIH-NCI P30 CA054174 (Cancer Center at UT Health San Antonio) and NIH Shared Instrument grant S10OD030311 (S10 grant to NovaSeq 6000 System), and CPRIT Core Facility Award (RP220662).

